# FAST: Filamentous Actin Segmentation Tool for quantifying cytoskeletal organization

**DOI:** 10.1101/2025.09.22.677898

**Authors:** Vineeth Aljapur, Adam Gardner, Jason Carayanniotis, Andrew R. Harris

**Affiliations:** Department of Mechanical and Aerospace Engineering, 1125 Colonel by Drive, Carleton University, Ottawa, ON, K1S 5B6; Artinus Consulting Inc., Ottawa, ON

**Keywords:** Actin, Cytoskeleton, Fluorescence Microscopy, Segmentation, Deep learning

## Abstract

Studying how actin filaments are assembled into different subcellular structures can provide insights into both physiological processes and the mechanisms of disease. However, quantifying the size, abundance, and organization of different classes of actin structure from optical microscopy data remains a challenge. To address this, we developed a deep learning based Filamentous Actin Segmentation Tool (FAST) to accurately and efficiently segment and quantify different classes of actin structure from phalloidin stained confocal microscopy images. We evaluated the performance of this tool to segment and quantify the abundance of different classes of actin structure in different cell lines and with dynamic changes in actin organization using lifeact-GFP during drug treatments. FAST enables quantification of different classes of actin structure from actin images alone, without the need for specific antibodies against proteins in different actin structures and hence can be a useful tool for researchers studying for studying actin related pathways involved in cell motility, cancer metastasis, and drug development.

**Summary Statement:** We developed Filamentous Actin Segmentation Tool (FAST), that leverages deep learning and antibody assisted labeling to segment and quantify actin structures from optical microscopy images.

## Introduction

The actin cytoskeleton plays a critical role in a range of cellular functions including driving cell motility, determining cell shape and mechanical properties, and separating daughter cells during cytokinesis, to name a few (Dominguez and Holmes, 2011; Fletcher and Mullins, 2010; Pollard, 2016). This broad range of functions are enabled through the assembly of filamentous actin (F-actin) into a variety of specialized structures, each with unique characteristics (Fletcher and Mullins, 2010; Michelot and Drubin, 2011). Common classes of actin structure include lamellipodial and lamellar networks, filopodia, stress fibers, and focal adhesions, which are defined by the morphology of the structure, the organization of actin filaments within the structure, and the presence of specific actin regulatory proteins (Desroches and Harris, 2024; Harris et al., 2018). Lamellipodial networks are broad, thin sheet-like extensions that are 0.1-0.2 μm thick and several microns wide. They play a critical role at the leading edge of migrating cells and drive cell motility, wound healing, and immune cell functions (Arce et al., 2023; Innocenti, 2018). Lamellipodia consist of branched actin filaments formed through activity of the Arp2/3 complex (Suraneni et al., 2012). Filopodia are finger-like protrusions that are 0.1-0.3 μm in diameter and range in length from a few microns to more than 35 µm long (Mattila and Lappalainen, 2008; Schäfer et al., 2011). They act as sensory structures and play an important role in exploratory cell migration (Davenport et al., 1993; Gupton and Gertler, 2007; Mattila and Lappalainen, 2008). Within filopodia, parallel bundles of actin filaments are crosslinked by the protein fascin with the motor protein myosin X located at filopodial tips (Gupton and Gertler, 2007; Yamashiro et al., 1998). Stress fibers are cytoplasmic bundles of actin filaments approximately 0.5-1 µm in diameter and range in length from several microns to tens of microns long (Burridge and Wittchen, 2013). One major function of stress fibers is to generate cellular contractile forces (Lu et al., 2008). Stress fibers contain anti-parallel actin filaments crosslinked by proteins such as α-Actinin, and the contractile activity of stress fibers is driven by Myosin II (Tojkander et al., 2012). Some stress fibers terminate at focal adhesion which are small, ∼1µm in diameter, structures located at sites of contact with the surrounding environment (Geiger et al., 2009; Kanchanawong et al., 2010). Focal adhesions contain a number of different proteins that include adhesion and adapter proteins such as Paxillin and Integrins (Askari et al., 2010; Riveline et al., 2001; Schaller, 2001).

Given that different actin structures fulfill specific functions, quantifying their presence, abundance, and localization can yield valuable information about the physiological state of a cell (Desroches and Harris, 2024). Fluorescence microscopy is the tool of choice for characterizing the organization of the actin cytoskeleton both in live cells, by expressing and imaging fluorescent fusion proteins to probes such as LifeAct (Riedl et al., 2008), f- tractin (Lopata et al., 2018), and actin binding domains (Burkel et al., 2007; Harris et al., 2020, 2019), and in fixed cells using fluorescently conjugated phalloidin (Adams and Pringle, 1991; Melak et al., 2017). Furthermore, actin staining with phalloidin can be combined with immunostaining for actin regulatory proteins to assist with identifying a particular class of actin structure. For instance, myosin X is involved in the initiation and extension of filopodia and localizes to filopodial tips. Immunostaining for Myosin X has therefore become a common approach for identifying filopodia in microscopy images. Critically, the efficacy of this approach is highly dependent on the signal to noise ratio and quality of image that can be obtained with a particular antibody to obtain reliable identification of different classes of actin structure. Another limitation arises from the availability and cross reactivity of antibodies for multiple different classes of actin structures that can be used in tandem, for multi-class identification (i.e. host species, limited number of fluorescence imaging channels).

Beyond simply identifying actin structures with antibodies against actin regulatory proteins, image analysis methods to segment and quantify the abundance and localization different classes of actin structure are continually being developed (Desroches and Harris, 2024). For filopodial quantification, tools like filoVision (Eddington et al., 2024), Filoquant (Jacquemet et al., 2019), and filopodyan (Urbančič et al., 2017) are used with phalloidin staining to identify F-actin and in some cases with filopodial markers like myosin X. These tools identify the cell boundary using either a deep learning model (filoVision) or contour detection in ImageJ which is subsequently used to detect filopodial protrusions based on morphology (Filoquant, filopodyan) or with a filopodial marker (filoVision). Another ImageJ algorithm FilamentSensor (Hauke et al., 2023) is suited for detection of stress fiber related features, along with machine learning based plugins like CSBDeep (Boothe et al., 2023). Focal adhesions have been detected with imaging tools like CellProfiler (Carpenter et al., 2006) and SFAlab (Mostert et al., 2023) with thresholding, edge detection, and deep learning (Mohamed et al., 2024). Despite these advances, a key challenge is that these tools are typically developed for specific cell types and single classes of actin structure which constrain their widespread use. To address these challenges, we developed a unique approach using antibody assisted labelling to create high quality ground truth annotations for detecting multiple classes of actin structures. This dataset was used to train a deep learning pipeline, which we refer to as Filamentous Actin Segmentation Tool (FAST), to quantify the abundance and properties of actin structure classes.

## Results

### Antibody-assisted annotation

We sought to develop an image analysis approach that could segment multiple classes of actin structure from a phalloidin (F-actin) image alone, eliminating the need for using multiple antibodies to identify and segment different classes of actin structure at the inference step. Deep learning is particularly well suited to this task and has seen increasing use in optical microscopy image classification and segmentation tasks for images of cells and subcellular structures (Metlek, 2024; Stringer et al., 2021; Melanthota et al., 2022). Training a deep learning algorithm to perform image segmentation requires a high-quality labelled dataset to ensure accurate results. To generate a labelled dataset that can be used to segment different classes of actin structure for model training, we used antibodies targeting actin regulatory proteins to fluorescently label and identify different classes of actin structure. The localization of these antibodies could then be used to guide the annotation of 4 distinct classes of actin structure in phalloidin images namely, lamellipodia and lamellar, filopodia, stress fibers, and focal adhesions. We screened through a selection of antibodies that could be used for labelling based on three criteria (Fig. S1). First, we excluded antibodies that failed to produce a significant signal to background ratio (actin structure to cytoplasmic or non-specific staining). This characterization was based on colocalization of the signal from the antibody with phalloidin (Fig. S1). For this purpose, we calculated Pearsons’s correlation of the normalized pixel intensities from the line scan, and antibodies corresponding to correlation values less than 0.5 were rejected. Second, since we want to label several classes of actin structure in the same image for multi-class segmentation, we selected antibodies based on their compatibility with one another (i.e. availability, host species). Third, to minimize fluorescence crosstalk between imaging channels, fluorescently labelled secondary antibodies were chosen to ensure minimal overlap between fluorophore excitation and emission spectra, enabling clear distinction between different classes of actin structure. Based on these selection criteria, F-actin was labelled with Phalloidin 647 Reagent (Abcam, ab176759), Paxillin (Novus Biologicals, AF4259-SP) with Alexa Fluor 405 Donkey Anti-Sheep secondary (ab175676) was used to label focal adhesions, Myosin X (Novus Biologicals, NBP2-88926) with Alexa Fluor 488 Goat Anti Rabbit secondary (ab150077) for annotating filopodia, stress fibers were annotated with and Myosin II (Sigma-Aldrich, M8064-25UL) with Alexa Fluor 568 Goat Anti-Rabbit secondary (ab175471). To generate training datasets, we fixed and stained NIH-3T3 fibroblast cells (materials and methods). Following immunostaining, cells were imaged using a 60x Nikon water-immersion objective with a spinning disk confocal microscope (Fig. 1A, B). We collected 149 multichannel images across three biological replicates (∼50 images per replicate).

**Figure 1:**
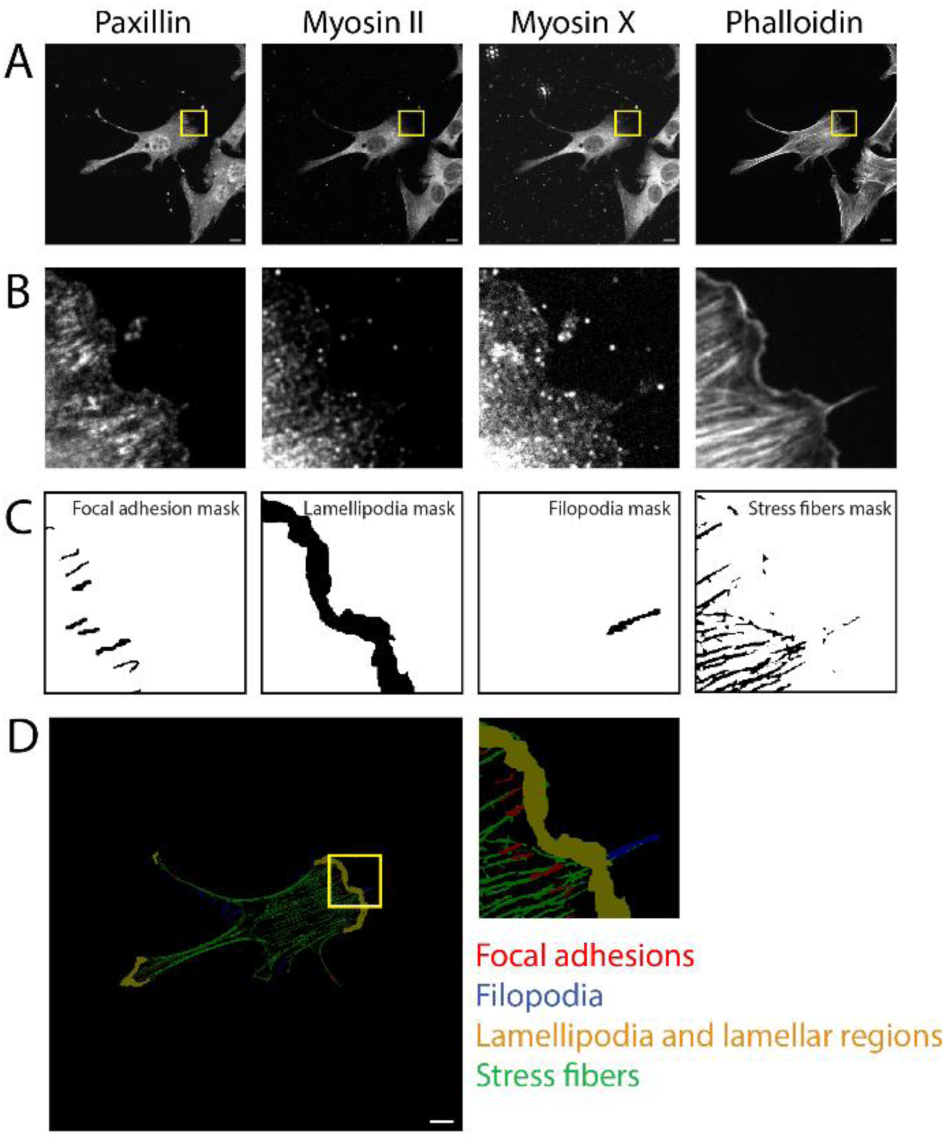
Antibody assisted labelling: **(A)** Representative images of a cell labelled with individual antibody channels of Paxillin, Myosin II, Myosin X, and Phalloidin. **(B)** Zoomed image of highlighted region (yellow box in A). **(C)** Focal Adhesions are detected with Paxillin channel, absence of Myosin II was used to identify lamellipodia and lamellar regions, presence of Myosin X puncta at the end of thin protrusions was used for detecting, and Phalloidin channel was used to detect stress fibers. **(D)** Composite mask (zoomed inset on right) from all individual annotations is used in training. All scale bars are 10 µm.

During data labelling, we ensured images contained only single cells by cropping centermost cell where applicable. To assist with annotation, on each of the above channels we performed background subtraction, contrast enhancement, and applied binary threshold prior to channel (Fig. S2). We used the following distinct visual signatures from image channels associated with each class of actin structure as a guide for semi-automated creation of annotation masks (Fig. 1A, B). To obtain Lamellipodial and lamellar regions we leveraged the channel corresponding to myosin II and phalloidin and highlighted the regions at the cell periphery where phalloidin is present, but myosin II is absent. Although this pattern is also applicable for filopodia, we confirmed filopodial presence by using the merged image of myosin X and phalloidin, as filopodial tips typically shown strong myosin X localization (Bohil et al., 2006). To detect stress fibers, we used images from the phalloidin channel alone. Note that even though there are multiple types of stress fibers (of dorsal, ventral, and transverse arcs) that are seen in our imaging, they were collectively treated as a single class of stress fibers for simplicity. For focal adhesions, we used images from the paxillin channel and highlighted the bright regions corresponding to focal adhesions on the cell. Stress fibers and focal adhesions were detected using the ridge detection and CSBDeep plugins of ImageJ respectively (Boothe et al., 2023). Masks were generated using a combination of automated and manual annotations, supervised by human-in-the loop verification (Fig. 1C). Manual annotation, verification, and correction of the masks were carried out using web-based annotation platform Supervisely (Supervisely, 2023). In total, 596 images were processed and annotated to create the training dataset.

### FAST training and evaluation

To train the model, the phalloidin channel for a single cell was used as the input image, with a mask containing four labels for different classes of actin structure. We used a supervised learning approach and selected an extension of UNet model with enhanced skip connections (UNet++) due to its fast inference speed and demonstrated success with biomedical datasets (Azad et al., 2024; Zhou et al., 2018). This model, equipped with a ResNet-34 encoder (He et al., 2016) (Fig. S3 A), was pre-trained on the ImageNet dataset (Deng et al., 2009). The model is configured to accept single-channel input images from phalloidin staining and produce an output segmentation mask with five classes, including background and four classes of actin structures of interest. To improve model generalization and reduce overfitting, we randomly transformed our dataset with added noise, cropped images, and altered brightness (Fig S3 B). The model was trained using dice loss for 100 epochs, with a dataset split of 99 training images, 20 validation images, and 30 test images, randomly selected across three experimental repeats (Fig S3 C, Methods: Training Parameters). Performance on validation images was used to select the model with the highest average dice score of 0.564.

To evaluate the model performance, we analyzed test images unseen during training and compared the annotated mask and predicted masks to get a comparable dice score of 0.569. A representative test image, along with the corresponding ground truth and predicted mask (Fig. 2A-C), as well as zoomed insets (Fig 2D-F) are shown. To assess whether there is any bias of detecting one class of actin structure over another, we calculated the fraction of each substructure and plotted the distribution of them against the same analysis from the ground truth masks (Fig. 2-G). All four substructures were found to be within the expected range between predicted fractions of cortical actin and stress fibers (0.73 ± 0.09), focal adhesions (0.06 ± 0.02), lamellipodia and lamellar regions (0.15 ± 0.08), and filopodia (0.07 ± 0.03) when compared with ground truth fractions of stress fibers (0.72 ± 0.07), focal adhesions (0.05 ± 0.03), lamellipodia and lamellar regions (0.17 ± 0.06), and filopodia (0.06 ± 0.03) with no statistically significant differences between the predicted and inference values for stress fibers (p = 0.67), focal adhesions (p = 0.37), lamellipodia and lamellar regions (p = 0.28), and filopodia (p = 0.48). In addition, OpenCV was used to count the number of detections for each actin class in the annotated masks and predicted masks. For stress fibers, we used line segment detector from OpenCV to count the fibers with actin label for ground truth and predicted masks, while the other substructures were counted with corresponding contours. These counts were plotted across all the test images (Fig 2-H) to assess the correlation between predicted counts and ground truth counts. A strong correlation was observed in filopodia (Pearson’s correlation = 0.84), stress fibers (Pearson’s correlation = 0.79), lamellipodia and lamellar regions (Pearson’s correlation = 0.61), and focal adhesions (Pearson’s correlation = 0.43). To assess experimental and biological variability we performed an analysis of random images that were reimaged under identical conditions (Fig. S4). No significant differences were observed between replicate imaging on the same cells after remounting the sample, indicating that our analysis tool was not influenced by the reproducibility of the imaging setup.

**Figure 2:**
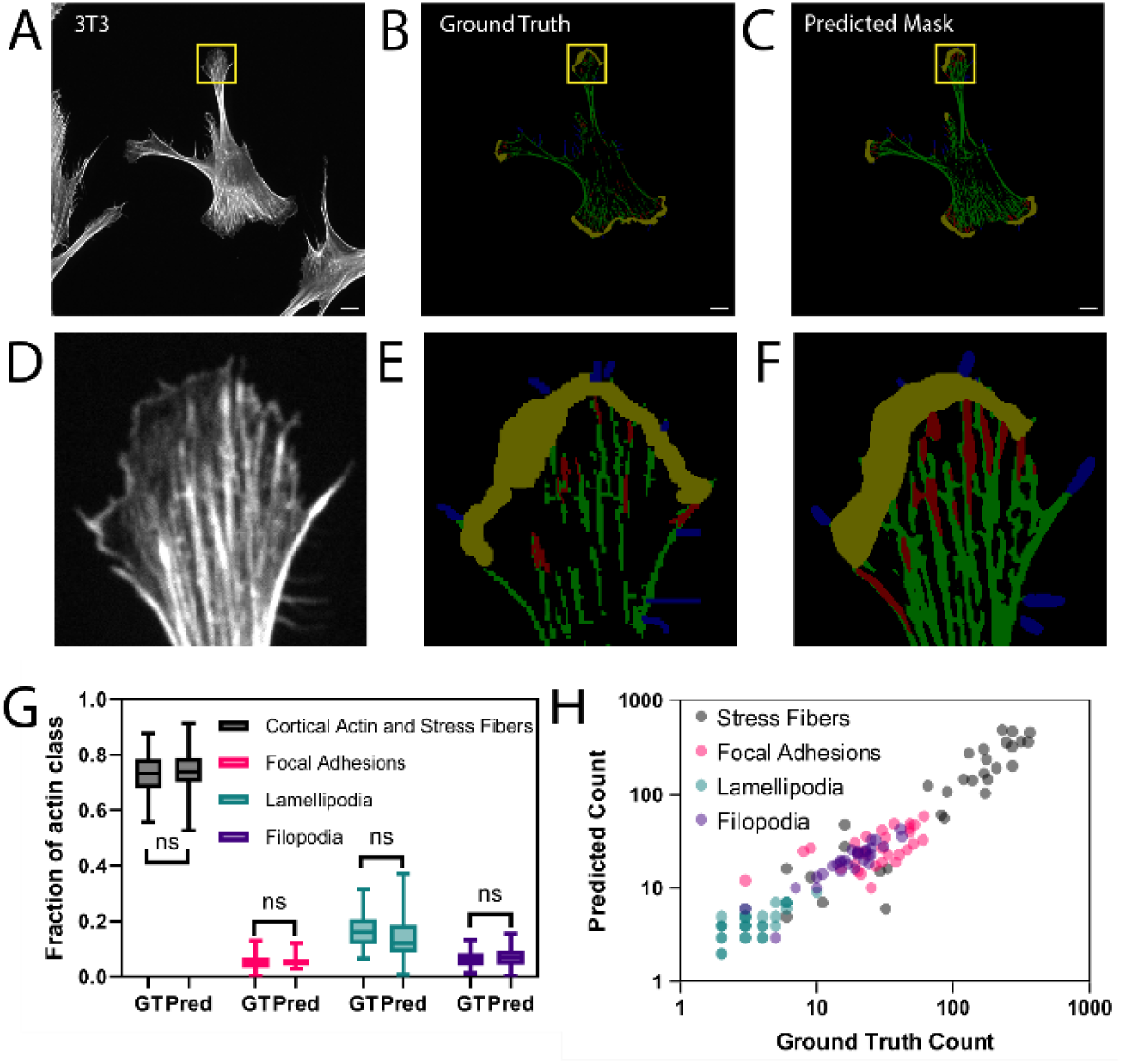
FAST inference evaluation for subcellar structure prediction: **(A)** Representative image of NIH-3T3 fibroblasts from Phalloidin channel used as input for algorithm testing. **(B)** Corresponding annotated mask with actin and stress fibers (green), focal adhesions (red), lamellipodia and lamellar regions (yellow), and filopodia (blue). **(C)** Corresponding predicted mask from the FAST model. **(D-F)** Magnified insets for highlighted regions of corresponding images. **(G)** For a random test set (n=30) the distribution of fractional area of each subtype from predicted masks is compared to that of the annotated mask to show no significant differences. **(H)** Scatter plot on log-log scale of the count of individual substructures from prediction mask for test set image and corresponding ground truth mask shown correlation between predicted counts with annotated counts. All scale bars are 10 µm.

### FAST analysis of different cell types

To evaluate the ability of FAST to accurately detect different classes of actin structure from images of different cell types, two commonly used epithelial cells lines were selected for validation: the human U-2 OS and the porcine LLC-PK1 (Methods: Cell Culture). Imaging was independently performed three times, following the same procedure used for NIH-3T3 fibroblasts, and ten images were collected from each session, resulting in 30 multichannel images per cell line. A representative image from each of the cell lines, along with its corresponding annotated and predicted mask, shown (Fig 3 A-F). In total, 240 images were processed and annotated with the same procedure previously described.

**Figure 3:**
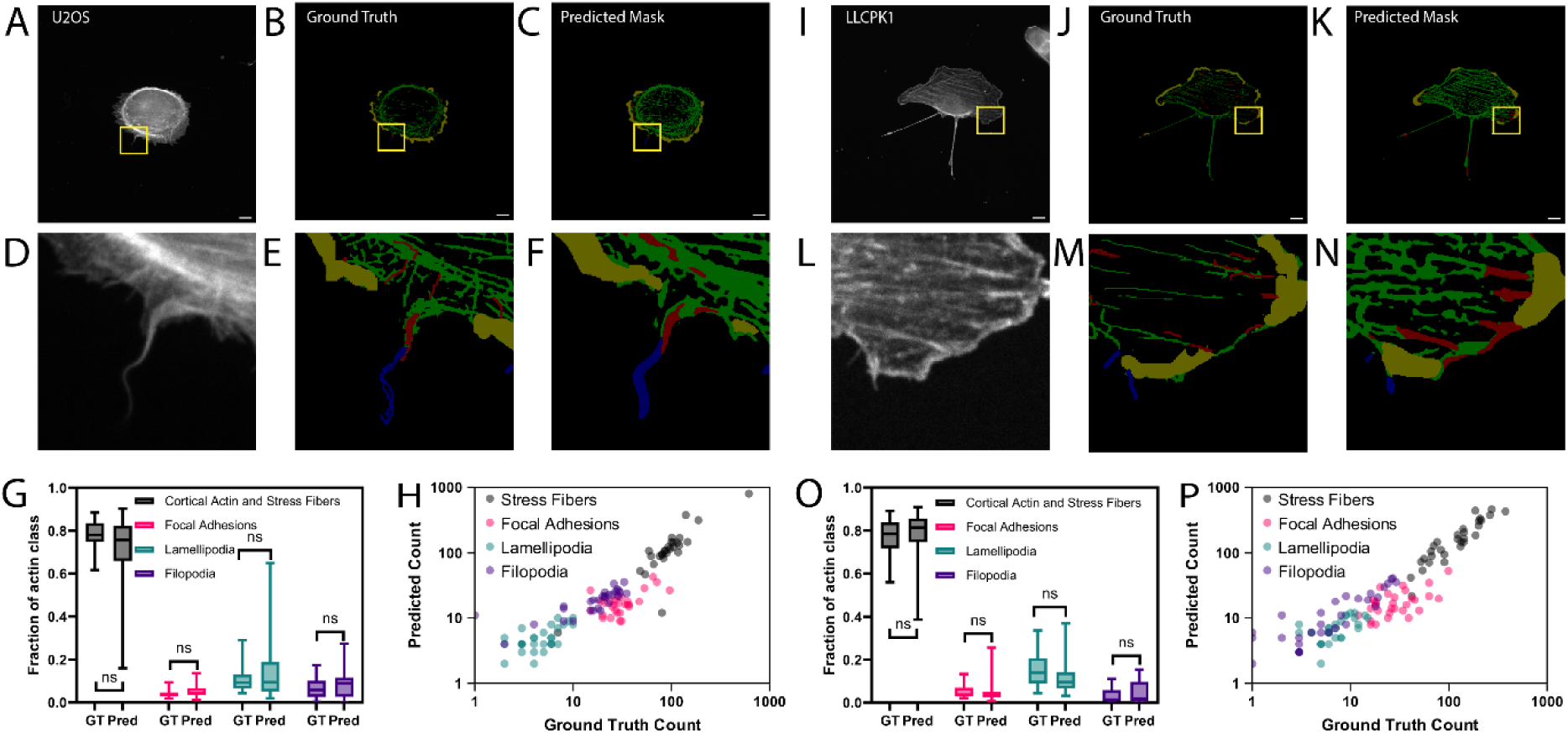
**FAST generalization evaluation on alternate cell types**: **(A)** Representative image from U-2 OS with **(B)** the ground truth, and **(C)** predicted mask. **(D-F)** zoomed insets of the images in **(A-C)**. **(G)** Comparison of the distribution of actin classes and **(H)** the number of each class and by counting the number of actin structures present in each image of U2OS. **(I)** Representative image from LLCPK1 with **(J)** the ground truth, and **(K)** predicted mask. **(L-N)** zoomed insets of the images in **(I-K)**. **(O)** Comparison of the distribution of actin classes and **(P)** the number of each class and by counting the number of actin structures present in each image of U2OS. All scale bars are 10 µm.

All collected images were processed with the trained model, which had not seen these images or these cell types during training. To assess model performance, we calculated the fraction of each class of actin structure and plotted their distributions (Fig 3G, I for U- 2 OS and LLC-PK1 respectively). The predicted fractions of U-2 OS for stress fibers (0.73 ± 0.15), focal adhesions (0.05 ± 0.03), lamellipodia and lamellar regions (0.13 ± 0.13), and filopodia (0.09 ± 0.07) closely matched the stress fibers (0.79 ± 0.07), focal adhesions (0.04 ± 0.02), lamellipodia and lamellar regions (0.11 ± 0.07), and filopodia (0.07 ± 0.05) of ground truth fractions with no statistically significant difference for stress fibers (p = 0.06), focal adhesions (p = 0.06), lamellipodia and lamellar regions (p = 0.33), and filopodia (p = 0.17) (Fig 5G). Similarly, the predicted fractions of LLCPK1 for stress fibers (0.79 ± 0.11), focal adhesions (0.05 ± 0.05), lamellipodia and lamellar regions (0.11 ± 0.07), and filopodia (0.05 ± 0.05) closely matched the cortical actin and stress fibers (0.77 ± 0.09), focal adhesions (0.05 ± 0.03), lamellipodia and lamellar regions (0.15 ± 0.07), and filopodia (0.03 ± 0.03) of ground truth fractions with no statistically significant differences between stress fibers (p = 0.4), focal adhesions (p = 0.95), lamellipodia and lamellar regions (p = 0.05), and filopodia (p = 0.16).

The ground truth mask and predicted mask were further analyzed by counting the number of each class of actin structure per cell (Fig. 3). This analysis on U-2 OS revealed a strong correlation in filopodia (Pearson’s correlation = 0.75), followed by stress fibers (Pearson’s correlation = 0.74), lamellipodia and lamellar regions (Pearson’s correlation = 0.5), and focal adhesions (Pearson’s correlation = 0.41). This was also seen LLC-PK1 cells with strong correlation in filopodia (Pearson’s correlation = 0.89), followed by stress fibers (Pearson’s correlation = 0.83), lamellipodia and lamellar regions (Pearson’s correlation = 0.49), and focal adhesions (Pearson’s correlation = 0.43).

### FAST analysis of live cell imaging data

To test the ability of FAST to quantify dynamic changes in the organization of the actin cytoskeleton in live cells, we expressed LifeAct-GFP in NIH 3T3 fibroblasts (Fig. 4A). Upon treatment of these cells with Rho-associated kinase (ROCK) inhibitor Y-27632 (50 μM), we observed notable differences in the abundance of different actin classes (Fig. 4B). For population analysis, we collected 90 images across three biological replicates (30 images per replicate) of cells treated with rock inhibitor Y-27632, and compared to 90 images across three biological replicates (30 images per replicate) of cells with vehicle control DMSO (Cytoskeletal Drug Treatments: Materials and Methods).

**Figure 4:**
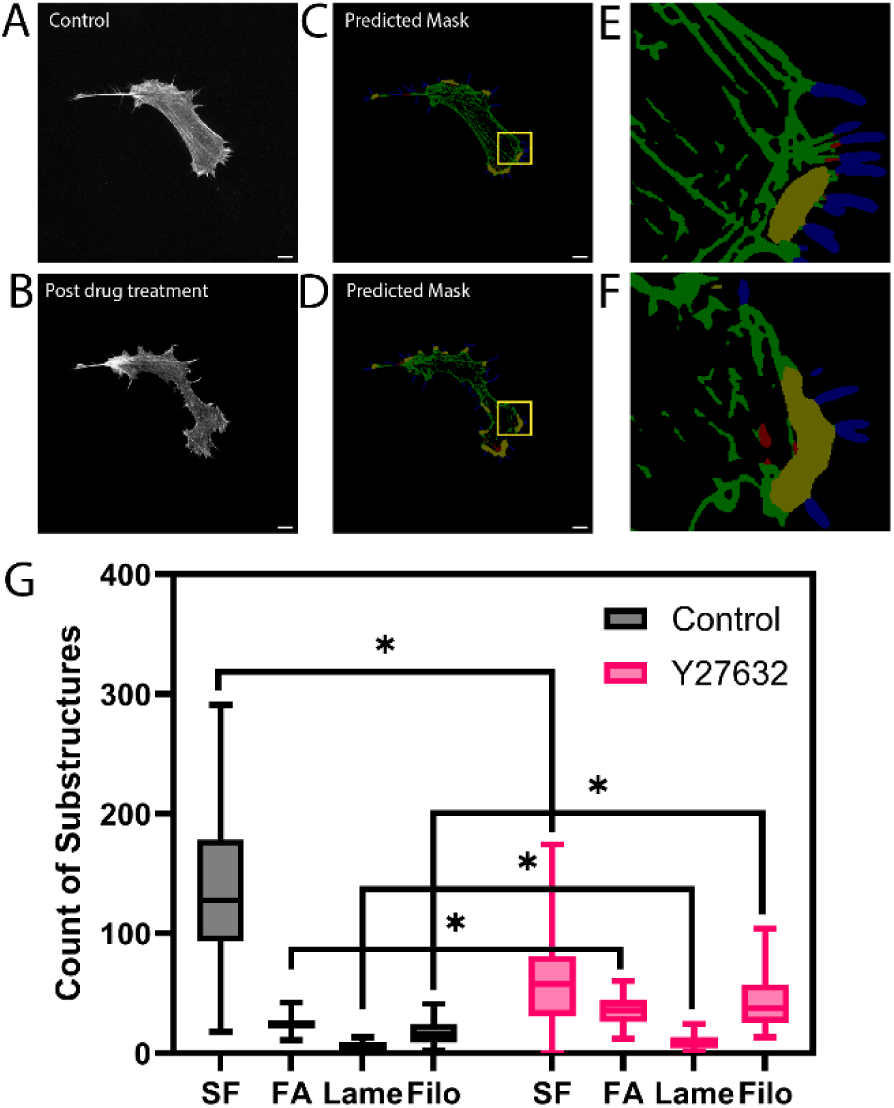
**FAST inference on live cell imaging data**: **(A)** NIH 3T3 fibroblasts expressing LifeAct-GFP. **(B)** Cells treated with the ROCK inhibitor Y27632. Predicted masks for a representative cell before **(C)** and after **(D)** drug treatment. **(E, F)** Zoomed in regions of corresponding predicted masks (highlighted in yellow box) **(G)** The count of classes of actin structure that are detected showing statistically significant changes in the abundance of different actin structures in treated cells. All scale bars are 10 µm.

For quantitative analysis, we counted the number of filopodia, lamellipodial and lamellar networks, stress fibers, and focal adhesions under each condition with OpenCV contour counts. Box plots of each label count showed significant differences among the drug conditions, especially there was a significant increase (p=1.3e-15) in filopodia count from control (18 ± 10) to rock inhibitor (43 ± 24) and lamellipodia count (p=7.4e-6) from control (6 ± 2) to rock inhibitor (9 ± 6) along with significant reduction (p=1.8e-19) in stress fibers from control (135 ± 56) following ROCK inhibition (61 ± 39).

## Discussion

In this work we developed FAST to enable accurate and automated detection of 4 different classes of actin structure using phalloidin images and live cell imaging with LifeAct-GFP expressed cells without needing additional antibodies. It can process thousands of images within an hour (more than 10 images per second) with reasonable sized GPUs and is well-suited for faster hardware acceleration with in-house and cloud computing if needed. The frozen model ensures consistent results for repeated runs of a given image free from human bias. Both the trained model and the labelled dataset are open-sourced (Data and resource availability), allowing researchers to use them as-is or fine-tune it further to meet specific needs.

Although this tool was trained with data augmentation to account for variation within data, it should be noted that the quality of the analysis largely depends on the quality of the image captured. Specifically, the signal to noise ratio for the image, extent of photobleaching, and the focus of the cell play a vital role in the accuracy of the analysis. While it is common practice to plate the cells at lower density to avoid overlapping cells and limit the number of cells per image, images containing multiple cells can be cropped prior to processing to analyze the cells individually.

To assess if our analysis could be applied to other cell types, we applied the trained model of FAST to datasets from different adherent cell types beyond the ones used in training. FAST was still able to identify the four classes of actin structures with reasonable accuracy, though statistical confidence was lower than with the original cell types. This suggests that while FAST can be used across a range of cell types, performance may benefit from finetuning on cell-type-specific data. Furthermore, fine-tuning can also help the model avoid misclassification of unseen classes of actin structure. For example, in our drug treatment experiments, disrupted F-actin was classified as focal adhesions resulting in higher focal adhesion count. However, these structures could more likely be actin puncta (Fig. 4G). In future work, further fine-tuning of FAST or the addition of other classes of actin structure such as cortical actin, puncta, asters and stars could improve the model performance and its general use for other cell types and treatments.

As there are multiple tools available for actin cytoskeleton analysis, the optimal choice of the tool ultimately depends on the volume of data and the types of antibodies or markers used. In cases where there are only a few images to be analyzed, manual approaches using tools like ImageJ might be sufficient. FAST is best suited to analyze large number of images for classes of actin structure from Phalloidin stained fixed cells and life cell imaging expressing signal for F-actin. However, if the dataset includes markers specific to a single substructure, alternative tools may also be adequate. For example, for single class quantification of filopodia with Myosin X antibody, filoVision could be a more suitable option.

To gain a comprehensive understanding of cellular processes, it is essential to analyze different classes of actin structure simultaneously- a feature lacking in most current tools. Additionally, these tools also need to be compatible with diverse cell types and live cell imaging to be widely useful. Unfortunately, many of the existing tools are not well suited to quantify dynamic changes in live cell imaging. This highlights the need for actin quantification tool like FAST, which can simultaneously detect multiple structures from diverse cell lines in dynamic environments like live cell imaging. Moreover, it is a resource and time consuming to source different antibodies to detect each kind of actin substructure individually. FAST was developed to address the need for detection of classes of actin structure without multiple specific antibodies, making them incompatible to quantify dynamic changes. Antibody assisted labelling approach has shown potential to train models for quantification of the actin cytoskeleton.

## Materials and methods

### Cell Culture

NIH-3T3 wild-type fibroblasts, U-2 OS (ATCC HTB-96 ™), and LLC-PK1 (ATCC CL-101 ™) cells were cultured in Dulbecco’s Modified Eagle’s Medium (DMEM) supplemented with 10% Fetal Bovine Serum (FBS) and 1% penicillin-streptomycin (PS) solution. Cultures were maintained at 37°C in 5% CO2 and passaged when confluent using 0.25% Trypsin-EDTA solution. For experiments conducted on 8-well imaging chambers (CellVis), the chambers were coated with 10 μg/ml fibronectin and incubated for 1 hour at 37°C. Then the cells were plated at a lower density and incubated for 6 hours in DMEM supplemented with FBS and PS.

### Antibodies and reagents

Primary antibodies used in this study include Paxillin antibody with sheep host (Novus Biologicals, AF4259-SP), Integrin beta 5 Antibody (Novus Biologicals, AF3824-SP), myosin X antibody with mouse host (Novus Biologicals, NBP2-88926), Anti-MYO10 antibody with rabbit host (Abcam, ab224120), Lamellipodin Antibody (H-5) with mouse host (Santa Cruz, sc-390050), Lamellipodin (D8A2K) with rabbit host (Cell Signaling Technology, 91138T), Anti-Myosin IIA (Sigma-Aldrich, M8064-25UL), Phospho-Myosin Light Chain 2 (Ser19) Antibody (Cell Signaling Technology, 3671T), Arp3 Antibody (A-1) Alexa Fluor® 488 (Santa Cruz, sc-48344 AF488), Anti-Paxillin antibody [Y113] with rabbit host (Abcam, ab32084), and Phalloidin-iFluor 647 Reagent (Abcam, ab176759) for staining actin. Secondary antibodies used in this study include Donkey Anti-Sheep IgG H&L (Alexa Fluor® 405) (Abcam, ab175676), Donkey Anti-Mouse IgG H&L (Alexa Fluor® 488) (Abcam, ab150105), Goat Anti-Rabbit IgG H&L (Alexa Fluor® 568) (Abcam, ab175471). Both primary and secondary antibodies were diluted in blocking solution according to the manufacturer’s protocol.

### Cytoskeletal Drug Treatments

Pharmacological treatments involving cytoskeletal drugs were used as agonists to form actin subtypes. To obtain microscopy images of cells that had undergone cytoskeletal drug treatments, the cells were incubated in 8-well chamber with DMEM supplemented with FBS and PS for 6 hours. 30 minutes prior to performing the immunostaining, the cytoskeletal drugs were added to DMEM at the following concentrations: 50 μM of Y- 27632 dihydrochloride, Rho kinase inhibitor (Abcam, ab120129). For control well the DMEM was mixed with 1.5 μl of Dimethyl sulfoxide (DMSO) (Invitrogen, D12345) as a vehicle control. The cells were then prepared for imaging according to the protocol described below under ‘Immunostaining’ for single channel.

### Immunostaining

Cells were fixed in 4 % paraformaldehyde in cell culture water supplemented with 10 % cytoskeleton buffer (10 mM PIPES/KOH, 100 mM NaCl, 300 mM sucrose, 1 mM EGTA, and 1 mM MgCl2 in distilled H2O) and 0.1 mg/mL sucrose for 20 min at 4 °C. Cells were then washed 3 times with Phosphate Buffered Saline (PBS), and permeabilized with 0.1 % Triton X-100 in PBS for 20 min at 4 °C. Cells were again washed 3 times with PBS and subsequently blocked with 2 mg/mL BSA in PBS overnight at 4 °C to prevent non-specific binding.

For single channel imaging experiments that do not involve actin subtype antibody binding (like drug treatment and filopodial count in cancer cell experiments), cells were then stained with Phalloidin-iFluor 647 Reagent (Abcam, ab176759) at a 1:200 dilution in 2 mg/mL BSA in PBS for 1 hr at 4 °C. Prior to imaging, the staining solution was removed from each well and rinsed twice with PBS and subsequently replaced with PBS containing 0.1% sodium azide (Sigma-Aldrich, 08591-1ML-F).

For multi-channel imaging experiments, after blocking the cells were then added with primary Paxillin antibody with sheep host (Novus Biologicals, AF4259-SP) at a 1:200 dilution in 2 mg/mL BSA in PBS for 3 hrs at 4 °C. This solution was removed from each well and rinsed 3 times with PBS and subsequently replaced with the secondary antibody Donkey-anti-Sheep IgG H&L (Alexa Fluor® 405) (Abcam, ab175676) at a 1:200 dilution in 2 mg/mL BSA in PBS for 1 hr at 4 °C. After which this solution was removed from each well and rinsed 3 times with PBS and subsequently replaced with the primary Anti-Myosin IIA (Sigma-Aldrich, M8064-25UL) at a 1:200 dilution in 2 mg/mL BSA in PBS for 3 hrs at 4 °C. This solution was removed from each well and rinsed 3 times with PBS and subsequently replaced with the secondary antibody Goat Anti-Rabbit IgG H&L (Alexa Fluor® 568) (Abcam, ab175471) at a 1:200 dilution in 2 mg/mL BSA in PBS for 1 hr at 4 °C. For next step, this solution was removed from each well and rinsed 3 times with PBS and subsequently replaced with the primary myosin X antibody with mouse host (Novus Biologicals, NBP2-88926) at a 1:200 dilution in 2 mg/mL BSA in PBS for overnight at 4 °C. This solution was removed from each well and rinsed 3 times with PBS and subsequently replaced with the secondary antibody Donkey Anti-Mouse IgG H&L (Alexa Fluor® 488) (Abcam, ab150105) at a 1:200 dilution in 2 mg/mL BSA in PBS for 1 hr at 4 °C. Finally, cells were then stained with Phalloidin-iFluor 647 Reagent (Abcam, ab176759) at a 1:200 dilution in 2 mg/mL BSA in PBS for 1 hr at 4 °C. Prior to imaging, the staining solution was removed from each well and rinsed twice with PBS and subsequently replaced with 0.1% Sodium azide/PBS

### Imaging Setup

Cells were imaged using Nikon Eclipse Ti2 inverted microscope utilizing a CrestOptics X-Light V3 spinning disk integrated with a Photometrics Kinetix camera with 60x Apo Plan water objective. Images were acquired using the NIS-Elements software v6.10.01 at a resolution of 1600 by 1600 pixels with pixel length of 0.108 µm. The exposure time for single channel imaging of phalloidin was set to be 200 ms with 5-10% laser power.

### Training Parameters

Full sized grayscale images (of resolution 1600x1600) were used by input images with a batch size of 3. The dataset is randomly split into training, validation, and test sets with 100-20-30 sizes. The UNet++ model was trained for 100 epochs on Nvidia A100 (40GB) with a learning rate of 0.001 using an Adam optimizer. Data augmentation was applied on each epoch by cropping (of resolution 800x800) at random locations, altering brightness (by factor between 0.5 to 1.5), and adding gaussian noise (mean=0.0, std=0.1) independently to roughly half of the images (p=0.5). The loss function used was dice which could be obtained from dice coefficient which for a given sets A and B defined as

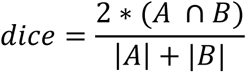

The best performing model on the validation set was selected and evaluated on test dataset as shown in the results.

### Software Tools Used

Data was preprocessed using ImageJ distribution of Fiji v2.16.0 (Schindelin et al., 2012) for removing noise and reducing cytoplasmic signal. The images were semi annotated using imageJ plugin CSBDeep (Boothe et al., 2023). Manual annotation of images was done using Supervisely (Supervisely, 2023) where each pixel was assigned a class value of zero to five where zero is (default) background, one corresponds to stress fibers and cortical actin, two corresponds to focal adhesions, three represents lamellipodia and lamellar regions, and four shows filopodial protrusions. Deep learning model was trained using PyTorch v2.6.0 on the annotated dataset. The training code along with a model inference pipeline was version controlled with GitHub and can be found in the data availability section. Plots were prepared using PlotNeuralNet v1.0.0, GraphPad Prism v10.2.2, and Adobe Illustrator v29.3.1.

### Statistical analysis

Data are presented as mean ± standard deviation. Unpaired two-tailed t-tests with Welch’s correction were performed to assess statistical significance between samples, and p-values of <0.05 were considered significant. Models were evaluated using the dice score (above) with 10-fold cross-validation, selecting the model with the best validation score.

## Author Contributions

All authors were involved in conceiving and designing the study. VA performed the experiments and analyzed the data. VA developed the deep learning model and collected training datasets. VA and AH drafted the manuscript. AG and JC provided feedback on model development and implementation.

---

**Figure S1.**
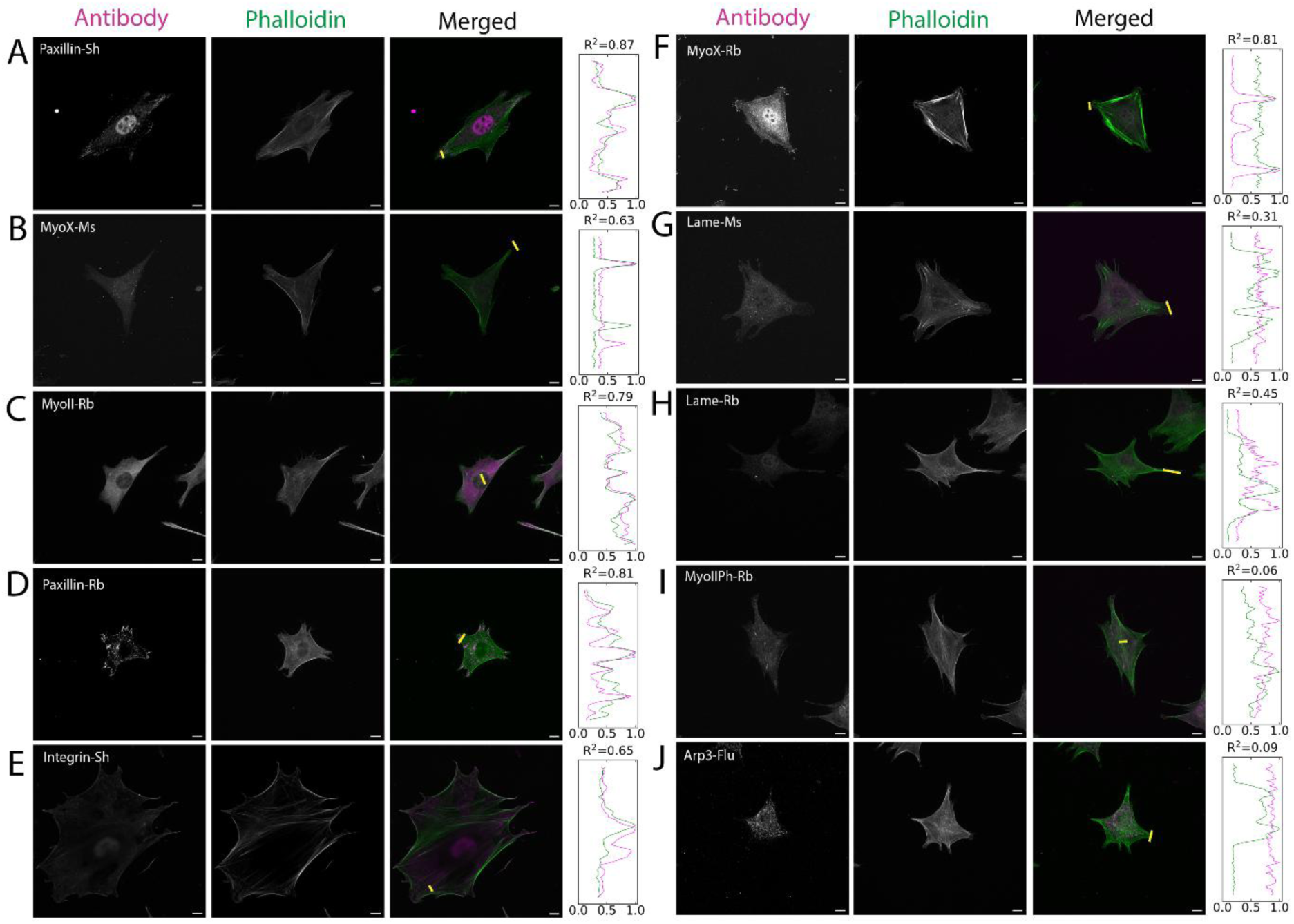
Overview of antibodies that were tested: channels with images (from left to right) corresponding to antibody channel, and phalloidin channel, and merged channel respectively. Pearson’s correlation (R^2^) was reported for each antibody based on the line scans of region corresponding to yellow line on the merged channel. **(A)** Paxillin antibody (AF4259-SP) with sheep host (R2=0.86), (**B)** Myosin X antibody (CL8994) with mouse host (R2=0.63) **(C)**, Myosin II antibody (M8064-25UL) with rabbit host (R2=0.79) were selected along with Phalloidin for this experiment. **(D)** Paxillin antibody (ab32084) with rabbit host (R2=0.8) showed promising results but was incompatible with the procedure due to common host. **(E)** Integrin antibody (AF3824-SP) with sheep host (R2=0.65) was effective but lost the specificity after couple of months of use. **(F)** myosin X antibody (ab224120) with rabbit host (R2=0.81) showed promising results but was incompatible with the procedure due to common host. **(G)** Lamellipodin antibodies with mouse host (sc-390050) (R2=0.31) and **(H)** rabbit host (91138T) host (R2=0.45) were ineffective in distinguishing lamellipodia. **(I)** Anti-Myosin Light Chain 2 antibody with rabbit (ab92721) host (R2=0.06) and **(J)** fluorescently conjugated Arp3 antibody (sc-48344 AF488) (R2=0.09) failed to obtain specific binding. All scale bars are 10µm.

**Figure S2.**
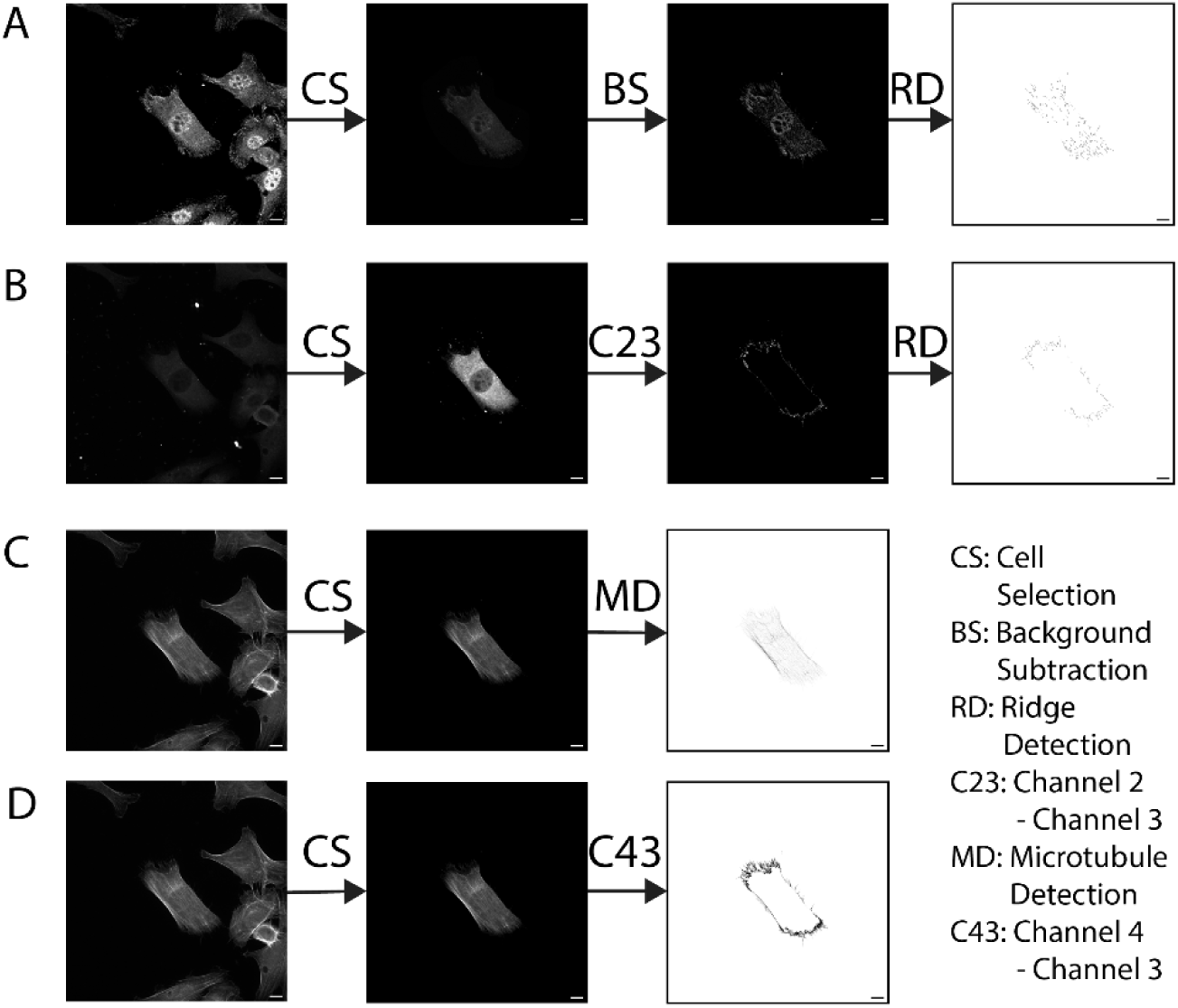
Data processing for mask generation: This involves splitting the multichannel image into individual channels, where each channel is associated with an antibody and processing them individually (from left to right) starting with cropping a single cell at the center. Background subtraction with rolling ball radius 50 was performed on paxillin channel **(A)** followed by ridge detection. For detecting the filopodia Myosin X channel **(B)** was subtracted with myosin II channel **(C)**. Lamellipodia was highlighted by subtracting Myosin II channel from Phalloidin channel **(D)**. For detecting stress fibers and cortical actin, background subtraction with rolling ball radius 20 was performed on phalloidin channel and the resulting image was run with CSBDeep plugin of Fiji/ImageJ. All scale bars are 10µm.

**Figure S3.**
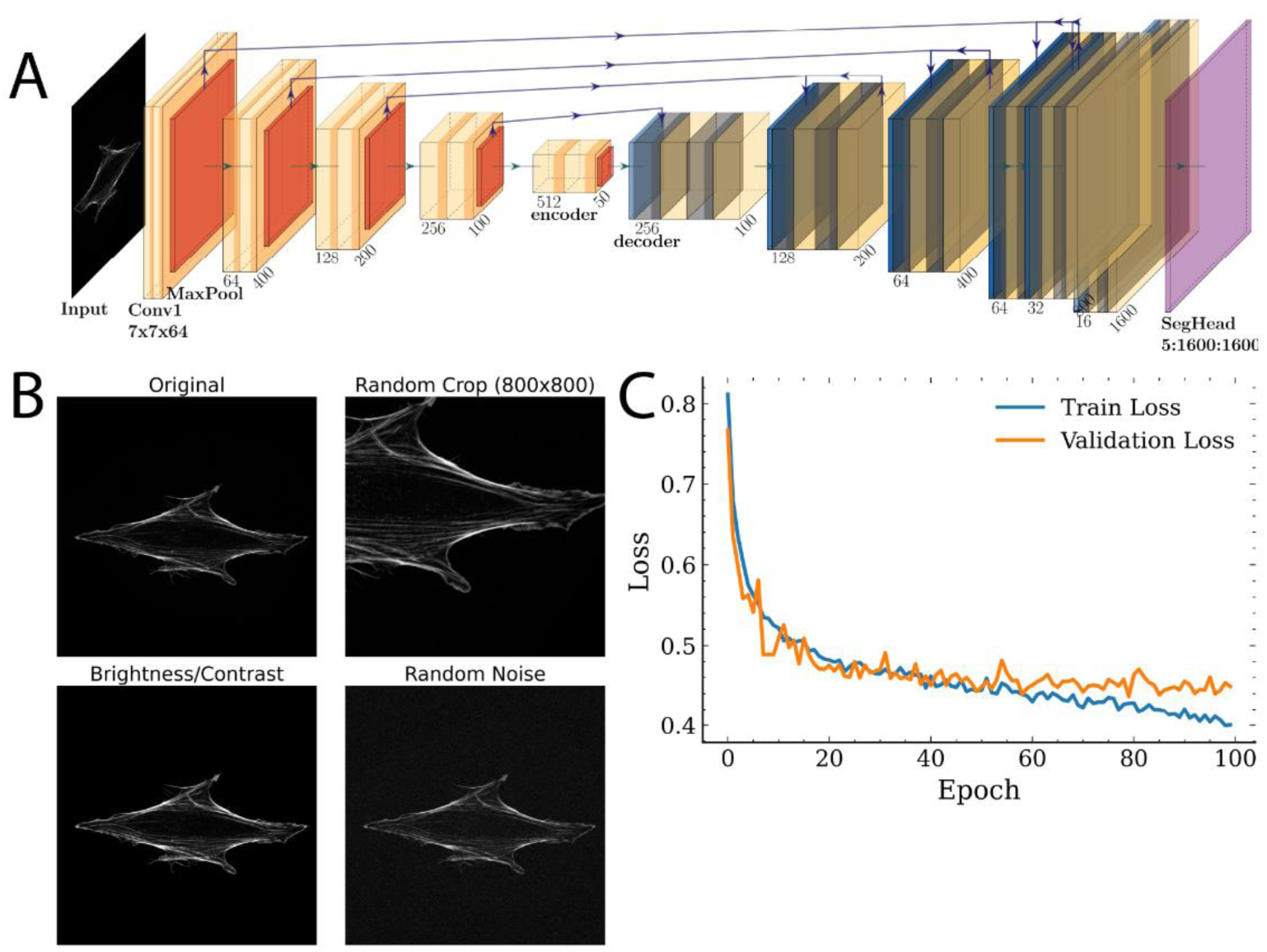
Model architecture and training with a schematic diagram: **(A)** Diagram of the network consisting of encoder and decoder with convolutional layers, pooling, skip connections for preserving information across the layers, followed by the final classification layer. The input on left is a grayscale image of 1600 by 1600 resolution and the output layer corresponds to class probabilities with the same shape. **(B)** Data augmentation of random crop altered brightness, and addition of noise was performed on random images for training the model. **(C)** Dice Loss function during the training indicates the flattening of validation loss around 50 epochs. Model snapshot corresponding to best validation loss (dice_score=0.56) is saved as the final model during the training. All scale bars are 10µm.

**Figure S4.**
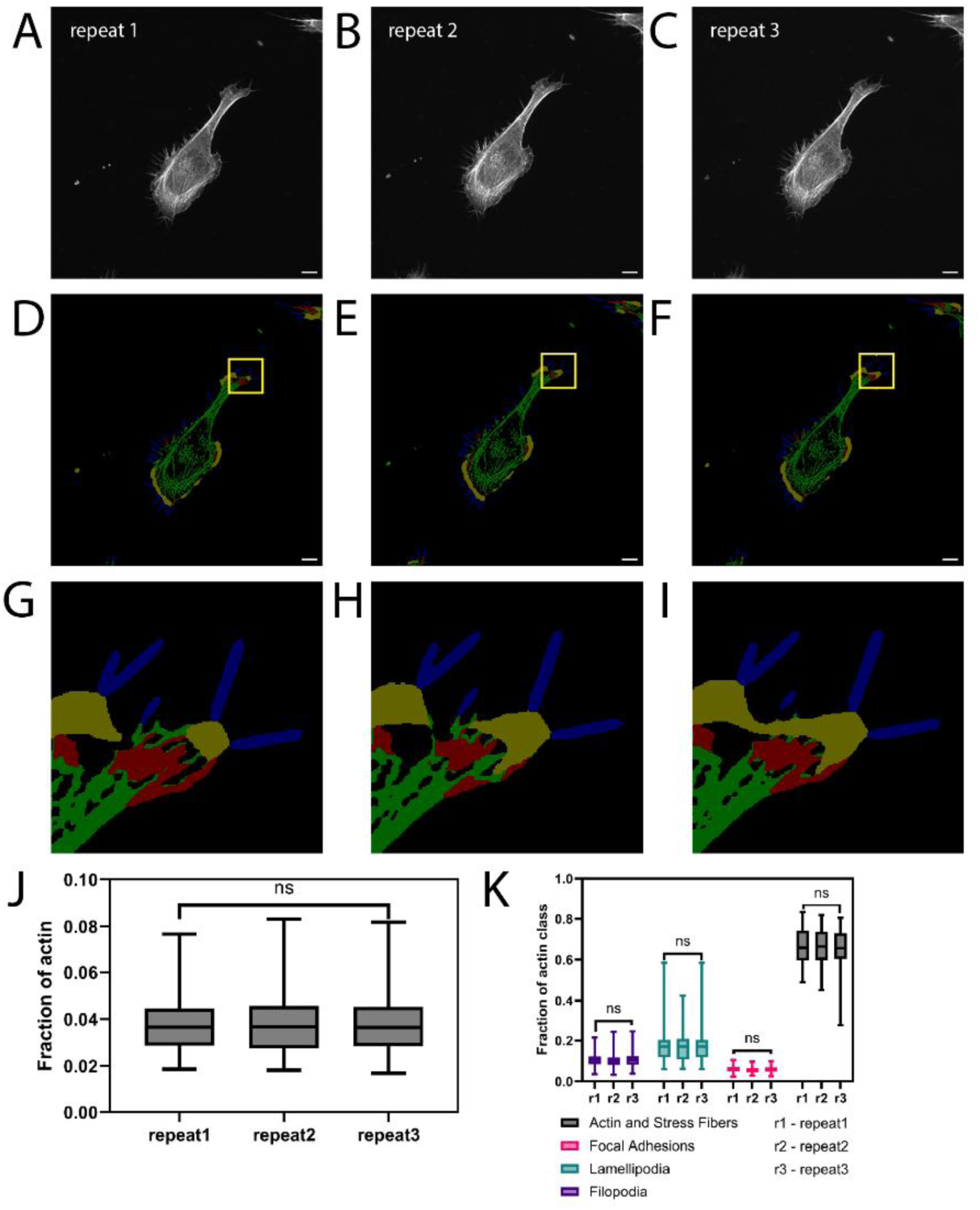
**Experimental and biological variability**: **(A)** Representative image from the random selection of images (n = 30) unseen during training that was assessed by model. To measure the variation across the experiments, images of these cells were recollected by finding the same cells **(B, C)** in subsequent repeats. **(D-F)** Predicted masks of corresponding representative image along with zoomed insets **(G-I)** (for region highlighted with yellow box) were shown. To estimate the loss in signal in subsequent repeats, the fraction of all pixels with phalloidin signal is estimated across the repeats **(J)** which was found to be insignificant. Biological variability is assessed by estimating the range of actin substructure distribution **(K)** across the selected cells for a given repeat. Subsequently, this distribution of classes of actin structure is compared by calculating the fraction of a given substructure for all the images across the repeats **(K)** to emphasize the differences in model predictions across the experiments. All scale bars are 10µm.

